# SOFisher: Reinforcement Learning-Guided Experiment Designs for Spatial Omics

**DOI:** 10.1101/2024.07.05.602236

**Authors:** Zhuo Li, Weiran Wu, Yan Cui, Jian Sun, Zhiyuan Yuan

## Abstract

Spatial omics technologies enable the precise detection of proteins and RNAs at high spatial resolution. Designing spatial omics experiments requires careful consideration of “what” targets to measure and “where” to position the field of views (FOVs). Current FOV sampling strategies often involve acquiring densely sampled FOVs and stitching them together, which is time-consuming, resource-intensive, and sometimes impossible. To optimize FOV sampling strategies, we developed SOFisher, a reinforcement learning-based framework that harnesses the knowledge gained from the sequence of previously sampled FOVs to guide the selection of the next FOV position, to improve the efficiency of capturing more regions of interest. We rigorously evaluated SOFisher’s performance using comprehensive simulations based on real spatial datasets, and our results clearly demonstrated that SOFisher consistently outperformed the conventional approach across various metrics. SOFisher’s robustness and generalizability were further validated through cross-domain generalization tests and its adaptability to varying FOV sizes. On a real Alzheimer’s Disease (AD) dataset, SOFisher successfully guided the selection of FOVs containing neurofibrillary tangles and amyloid-β plaques in both single and dual target tissue landmark scenarios. Remarkably, SOFisher-guided experiment design of spatial single-omics on limited tissue areas yielded insights into AD-related cell states, subtypes, and gene programs previously obtained through extensive spatial multi-omics experiments. SOFisher has the potential to revolutionize the experiment design of spatial biology.

## Introduction

Spatial omics technologies have revolutionized the study of tissue spatial biology^1^. These cutting-edge technologies can accurately detect dozens of proteins and hundreds of RNAs at a cellular spatial resolution, enabling in-depth spatial characterization of both healthy and diseased tissues^2,3^. To address specific research questions, scientists must carefully design their experiments.

In high-resolution spatial omics technologies such as imaging mass cytometry (IMC)^4^, multiplexed ion beam imaging by time of flight (MIBI-TOF)^5^, Co-Detection by Indexing (CODEX)^6^, Time-of-flight secondary ion mass spectrometry (TOF-SIMS)^7^, and in situ hybridization methods like seqFISH^8^ and MERFISH^9^, as well as the CosMx Spatial Molecular Imager (SMI)^10^, two of the most critical considerations in experiment design are (1) “what”: determining the set of targets to be measured (e.g., genes, proteins), which has been addressed extensively^11–16^, and (2) “where”: determining the positions of field of views (FOVs), which has not been adequately addressed. The standard approach to sampling a series of FOVs that cover regions of interest (ROIs) within a tissue as much as possible is to acquire densely sampled FOVs and stitch them together to cover a larger tissue area^17–19^. However, this process is time-consuming and resource-intensive, and certain techniques, such as TOF-SIMS and MIBI-TOF, may destroy FOV adjacent areas, making it impossible to stitch them together to form a complete tissue area. Furthermore, fully covering the tissue with densely sampled FOVs always leads to many unnecessary FOVs irrelevant to target questions, particularly in cases where the focus is only on specific regions containing target tissue landmarks (TTLs). For example, in Alzheimer’s disease (AD) research, scientists often focus on the niche surrounding amyloid-β plaques (Aβ) and neurofibrillary tangles (p-tau)^20,21^, while in liver research, the focus is often on the niche around landmarks such as central veins and portal nodes^22,23^. These factors underscore the significance of FOV sampling strategies in spatial omics experiments.

To circumvent the need for covering entire tissue areas, two studies have shifted attention to designing FOV sampling strategies^24,25^. They conducted statistical power analysis to study how factors like the number of FOVs is needed to maximize the detection of cell phenotypes. Since one only have very limited information about a tissue before the tissue is actually measured, the fundamentals of these previous studies are to position FOVs randomly (without replacement) and lack guidance of each FOV’s selection to enhance the likelihood of hitting the target. In fact, molecular and spatial phenotype information observed within each sampled FOV has the potential to offer valuable insights into its relative position within the tissue environment. This information could serve as a useful hint to guide the placement of the next FOV towards a higher likelihood of capturing TTLs. As the number of sampled FOVs increases, the sequence of sampled FOVs collectively contributes to a growing understanding of the entire tissue, providing informative cues for determining the optimal position of the next sampling FOV.

To enable a “smart” selection of FOV positions, we developed SOFisher, a reinforcement learning-based framework that optimizes sampling strategies for spatial omics experiments. SOFisher leverages knowledge from previously sampled FOVs to guide the selection of the next FOV position, ultimately improving the efficiency of capturing more ROIs. We designed comprehensive simulation datasets based on real spatial datasets to evaluate SOFisher’s performance, and SOFisher outperformed the baseline strategy in various metrics. We assessed the generalizability of SOFisher by training policies on several different aging stages of a mouse brain dataset and evaluating their performance on other different aging stages. SOFisher exhibited generalizability even with a domain gap between the training and testing data. Moreover, we demonstrated SOFisher’s compatibility with different FOV sizes to prove its adaptability to various real-world experimental settings. When applied to a real Alzheimer’s Disease (AD) dataset, SOFisher successfully guided the selection of FOVs containing both AD pathology markers of Aβ and p-tau. We demonstrated the practicability of SOFisher by using a SOFisher-designed spatial single-omics experiment on a small area to uncover key differential abundance cells, sub-types, and gene programs related to AD pathology, which was only possible by previous spatial multi-omics experiments on large areas. SOFisher simultaneously addressed three key gaps in current spatial omics experiment designs: the expense of whole tissue coverage, the challenge of FOV selection with limited knowledge of the tissue slice to be measured, and the infeasibility of single-cell spatial multi-omics technologies.

## Results

### Framework

When designing spatial omics experiments, scientists aim to sample multiple fields of view (FOVs) to cover the region of interest containing target tissue landmarks (TTLs). An optimal sampling strategy would ensure that every FOV contains TTLs, thereby enhancing the analytical power of associations between TTL-surrounding niches, cell states, cell-cell proximities, and microenvironments. However, it is almost impossible to obtain an optimal sampling strategy due to very limited knowledge (only the overall tissue boundary is known) about the tissue before the actual experiment. Consequently, the current design strategy can only perform random sampling within the slice boundary^24,25^. Furthermore, due to the infeasibility of single-cell spatial multi-omics technologies^26^, TTLs often cannot be profiled together with the spatial omics experiment itself, as they may belong to different layers of molecules (e.g., TTLs may be certain protein markers, while the spatial omics experiment may focus on measuring spatial gene expressions), resulting in the inability to observe TTL positions throughout the whole experimental process.

SOFisher (Fig. 1) aims to improve sampling efficiency beyond the currently utilized random sampling strategy by increasing the expectation of sampling more TTL-containing FOVs in a FOV-sampling sequence. The rationale behind SOFisher is that during the sampling procedure, each previously sampled FOV contains observations of spatially resolved cellular phenotypes in a small region of the tissue measured by spatial omics. By leveraging the knowledge obtained from all previous FOV sequences, SOFisher can guide the selection of the next FOV position to be sampled. SOFisher assumes that, regarding the cellular phenotypes observed by the spatial omics experiment, (1) the tissue is not randomly organized by cellular phenotypes, and (2) the TTLs have associations with the cellular phenotypes.

**Fig 1.**
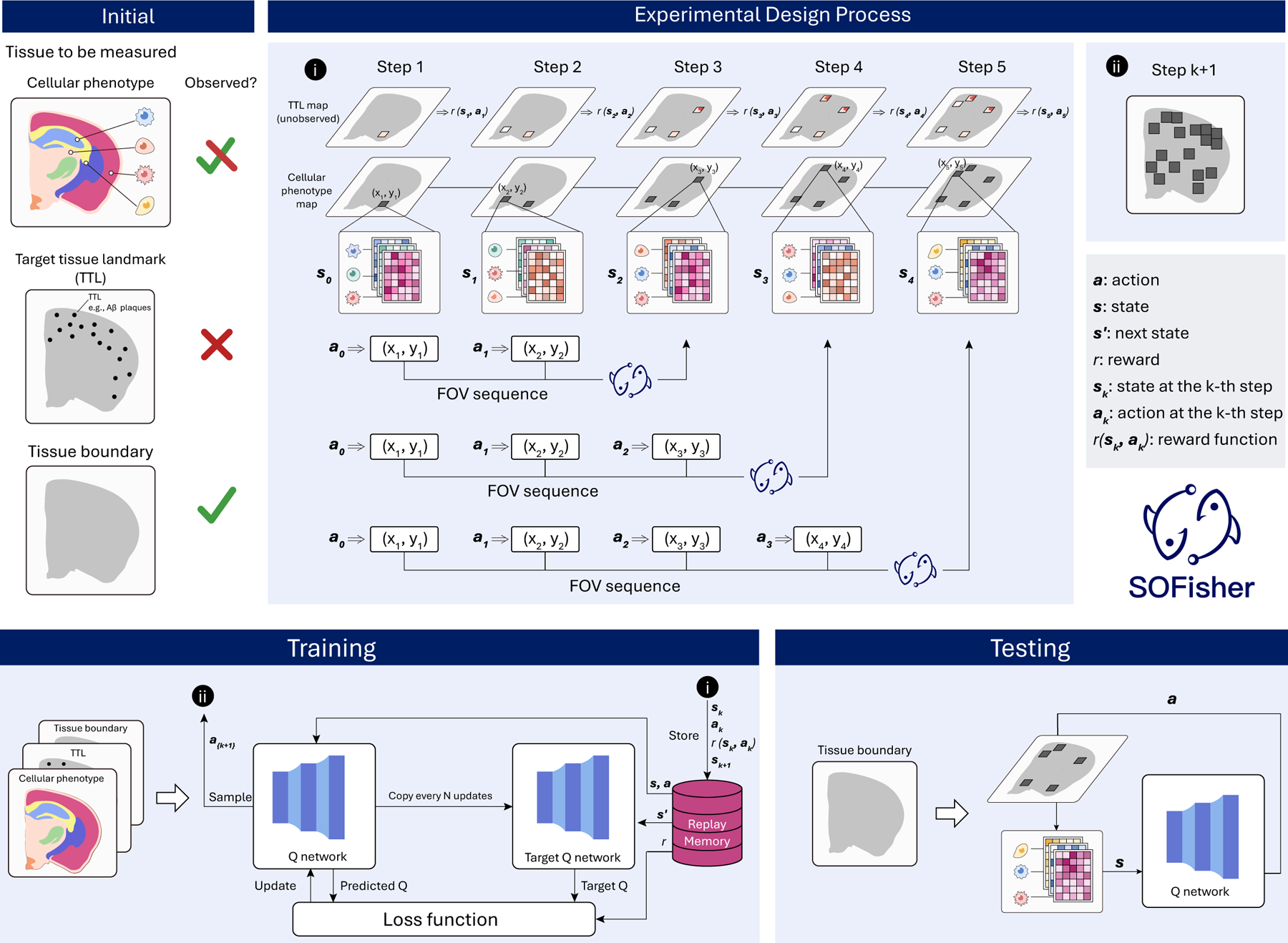
Workflow of SOFisher. Initial: Initial conditions of tissues to be measured in practice and in testing, i.e., observable cellular phenotype only in FOVs, unobservable TTL, and observable tissue boundary. Experimental design process: (i) An FOV sequence includes a number of steps. At the time step *k*, an action *a_k_* is selected according to the current SOFisher policy with state *s_k_*, which determines the sampling position in the tissue (*x_k+1_*, *y_k+1_*). By executing the sampling, a new state *s_k+1_* and the reward *r(s_k,_a_k_*) can be obtained. Training: The tuple *(s_k,_a_k,_r(s_k,_a_k_*), *s_k_*+1) at time step *k* from (i) is stored in a replay memory. At each step, a tuple (*s,a,r,s*’) is randomly selected for updating Q-network, where the objective is to minimize the loss function between the predicted Q from Q-network and Target Q from Target Q-network. In particular, Target Q-network copies parameters of Q-network every N update steps. At the next time step k + 1, the action *a_k + 1_* is sampled from Q-network with state *s*_k+1_ in (ii). Testing: Using the well-trained Q-network, an action α is output by the input of state *s* at each time step, where only the tissue boundary is observed.

During the training stage (Fig. 1), the training data contains both TTL and cellular phenotype information. Thus, SOFisher trains a policy that takes observations of previously sampled FOVs as input (e.g., cell types within FOVs) and selects the position of the next FOV by rewarding informative sampling steps (i.e., TTL-containing FOVs). The TTL information is only used for rewarding but not included in observations. At the testing stage (Fig. 1), the next FOV position is selected based on the observations in the previous FOVs according to the trained policy. TTL information is not observed throughout the testing process, the cellular phenotypes are only observable within sampled FOVs, and only the tissue boundary can be observed. The process terminates when the maximum number of FOVs is reached. Details can be found in Methods.

### Sampling Strategy for Target Tissue Landmarks with Cell Types Associations

We designed the first simulation dataset to evaluate SOFisher’s performance in sampling target tissue landmarks (TTLs) associated with cell types. The simulation was based on a real spatial transcriptomics dataset^17^ of the mouse primary motor cortex, measuring 258 spatially resolved gene expressions of approximately 300,000 cells on 64 slices from 2 mice (Fig. 2A). We generated TTLs around L45 IT cells with a probability of 20% to simulate TTL-cell-type association (Fig. 2B). At the beginning of the SOFisher sampling process, only the overall tissue boundary is known to SOFisher. During the sampling process, only cell type information within previously sampled FOVs can be observed, while cell types distributions in unmeasured FOVs and any TTL information remain unobserved. Running SOFisher on a single slice as an example (Fig. 2C), the sampling sequence on the cell type map and the TTL map are shown (Fig. 2D). Details of training and testing configurations can be found in Methods.

**Fig. 2.**
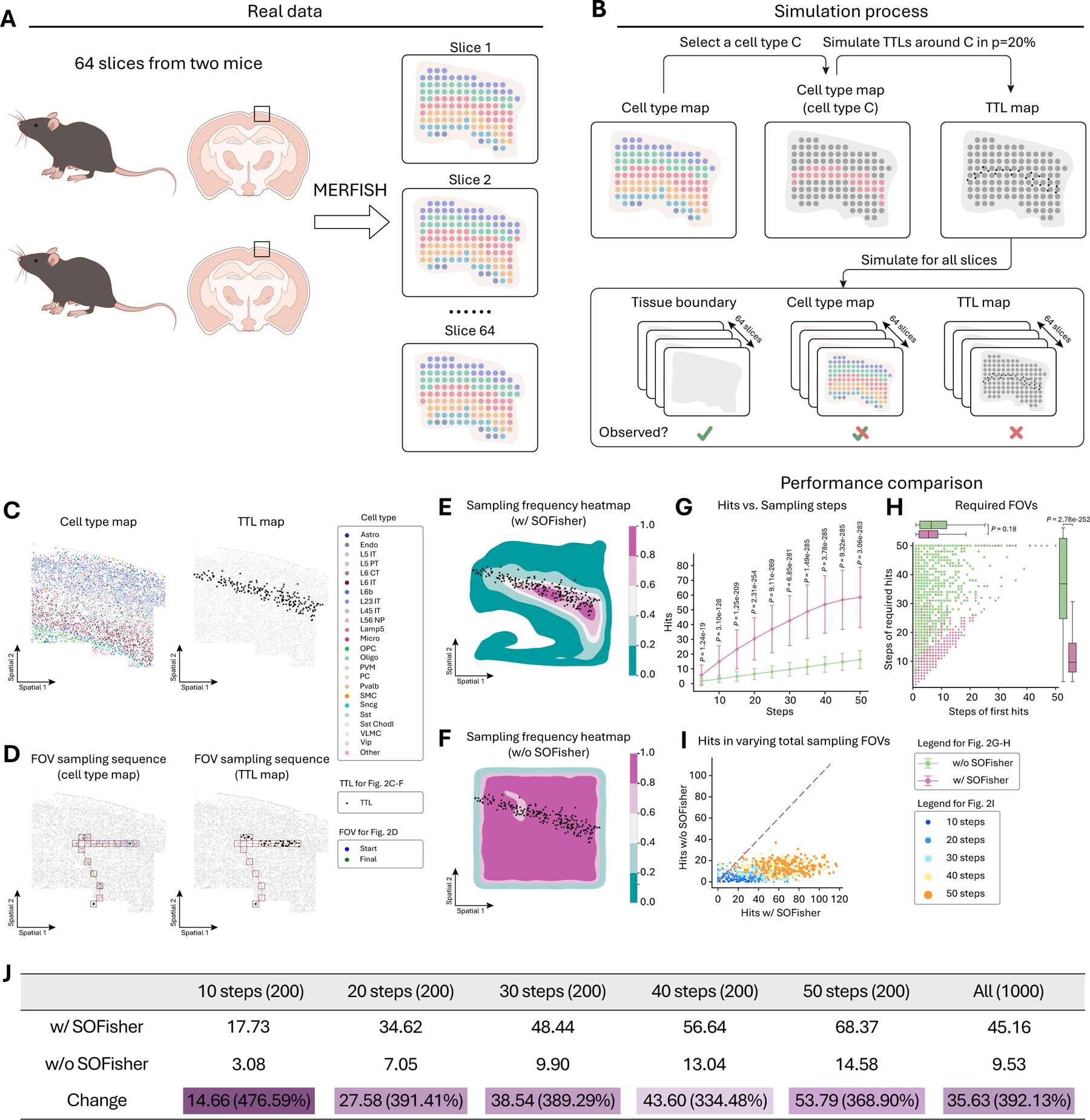
Sampling Strategy for Target Tissue Landmarks with Cell Types Associations. A: The procedure to obtain real spatial transcriptomics data of 64 slices from two mice by the spatial omics technology, MERFISH. B: The simulation procedure for TTLs: generating a TTL with probability 20% (p=20%) around each cell of the selected cell type C. Simulate the generation of TTLs for all 64 slices, and we can obtain 64 TTL maps. In particular, while the tissue boundary is totally observed, we can observe partial cell type map within FOVs and the TTL maps are not available. C: An example for cell type and TTL maps of a slice. In particular, we extend each slice into a rectangle and arbitrarily select one corner of the rectangle as the origin of a Cartesian coordinate system, with the two sides of the angle being the coordinate axes: Spatial 1 and Spatial 2. D: Example of the cell type and the TTL maps in an FOV sampling sequence in color with unsampled part in grey. E: The Sampling frequency heatmap on a slice with SOFisher policy (w/ SOFisher) for method evaluation, where we record the number of times each position on the slice is sampled within 1000 trails with randomly initial FOVs. F: The Sampling frequency heatmap on a slice with random policy (w/o SOFisher) for performance comparison. G: The variation in the number of TTL-containing FOVs (hits) with the increase of the total number of FOVs (steps) for the two compared policies of w/ and w/o SOFisher. The p-values (denoted by P) between the hits of the two policies at steps 5, 10, 15, 20, 25, 30, 35, 40, 45, 50. H: The minimum number of FOVs for obtaining a target number (=10) of hits, and the number of the first hit in a sampling sequence for 1000 trails with randomly initial FOVs. The p-values of the two performance indices between w/ and w/o SOFisher are given. I: Comparison in terms of the number of hits between w/ and w/o SOFisher within 10, 20, 30, 40, 50 steps. J: The second and third lines show the average of the number of hits over 200 trails within each of 10, 20, 30, 40, 50 steps and over the total 1000 trails, and the last line show the improvements of w/ SOFisher over w/o SOFisher.

To visualize the region-coverage likelihood of SOFisher, we repeated the sampling process 1000 times and recorded the sampled frequency of each cell, plotting the SOFisher-frequency heatmap on the TTL map (Fig. 2E). The results showed that the hotspots of the SOFisher-frequency heatmap largely overlapped with TTL localizations (Fig. 2E), in contrast with the random sampling strategy (Fig. 2F). We then compared the number of hits (i.e., TTL-containing FOVs) of different sampling strategies as a function of total sampling steps, indicating consistently better performance (up to 5 times improvement in sampling efficiency) of SOFisher compared to the random sampling strategy (Fig. 2G). Additionally, with approximately equal required steps for the first hit (i.e., the first step that sampled a TTL-containing FOV), the required steps for target hits (i.e., the number of FOVs that need to be sampled to obtain the target number TTL-containing FOVs) using SOFisher was significantly smaller than random sampling (Fig. 2H). Finally, we compared the performance of SOFisher and random sampling by independently performing the whole sampling process for 200 times and recording the number of hits for each time across different total sampling steps (Fig. 2I, J). The results showed that SOFisher outperformed random sampling in almost all cases, especially with a large number of sampling steps, indicating that more sampled FOVs can provide more knowledge about the tissue learned by the SOFisher policy (Fig. 2I, J). SOFisher’s improved performance was validated on another simulation data with different cell type-TTL associations (Supplementary Fig. 1).

### Sampling Strategy for Target Tissue Landmarks with Spatial Domain Associations

We designed the second simulation dataset to evaluate SOFisher’s performance in sampling TTLs associated with spatial domains. Unlike the first simulation dataset, where the TTLs were directly associated with the spatial omics observations (i.e., cell types), this dataset presents a more challenging scenario, as the TTLs are indirectly associated with the spatial omics observations through spatial domains. The simulation was based on a real spatial transcriptomics dataset^27^ of the mouse frontal cortex and striatum region (Fig. 3A). We generated TTLs around cells within the corpus callosum domain with a probability of 30% to simulate TTL-spatial domain association (Fig. 3B). Similar to the previous analysis, we ran SOFisher on a single slice as an example (Fig. 3C) and visualized the sampling sequence on the cell type map and the TTL map (Fig. 3D). By repeating the sampling process 1000 times, we generated the SOFisher-frequency heatmap, which largely overlapped with TTL localizations (Fig. 3E), in contrast to the random sampling strategy (Fig. 3F). Similar to previous analyses, SOFisher consistently outperformed the baseline method in some key metrics. First, SOFisher achieved a higher number of hits as a function of total sampling steps (Fig. 3G), indicating more efficient sampling of TTLs. Second, SOFisher required fewer steps to reach the target number of hits (Fig. 3H), demonstrating faster convergence to the desired sampling goal. Finally, for the same number of FOV samples, SOFisher obtained a higher number of hits (Fig. 3I, J), highlighting its superior performance in identifying TTLs. SOFisher’s improved performance was validated on another simulation data with different spatial domain-TTL associations (Supplementary Fig. 2).

**Fig. 3.**
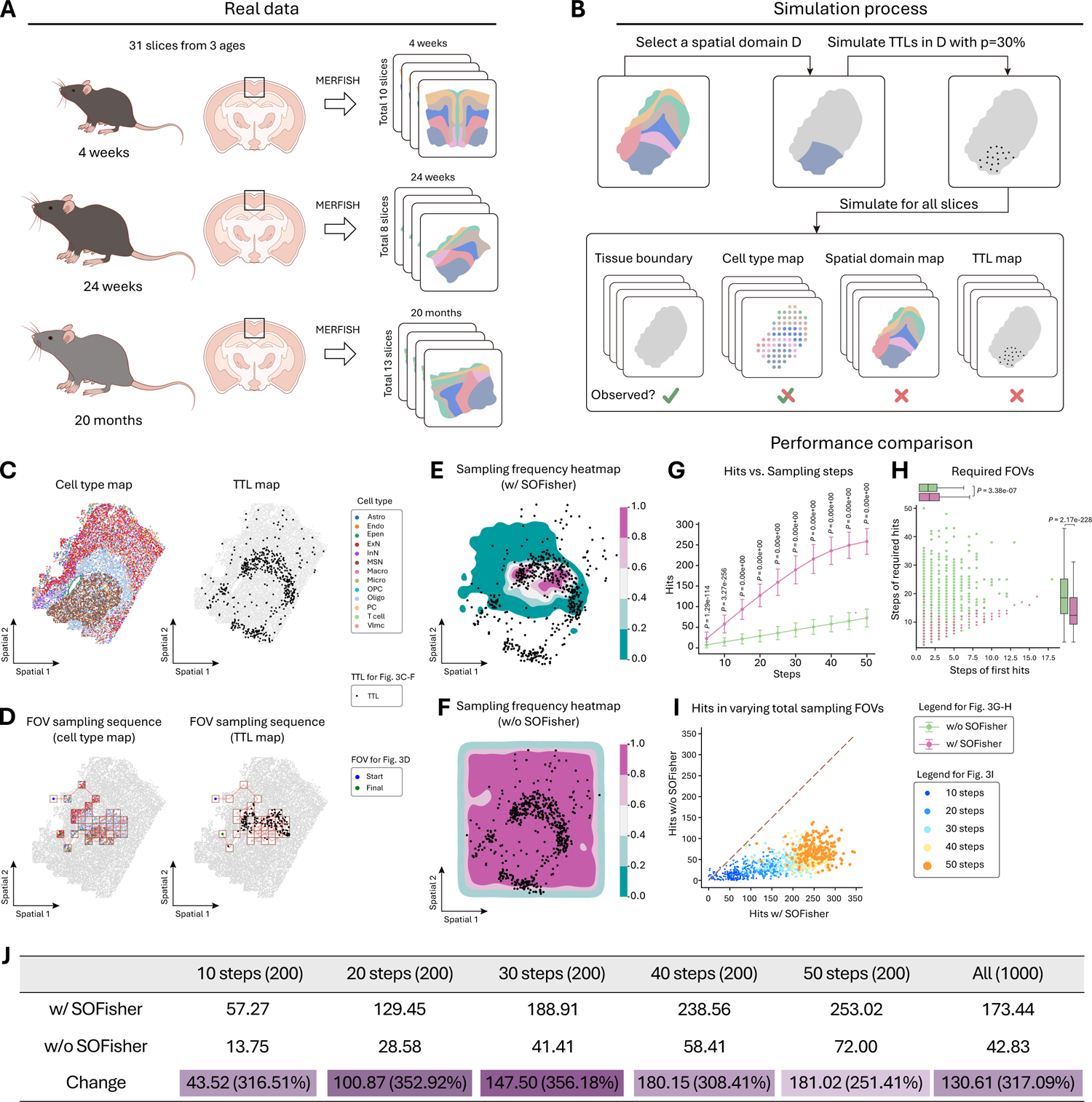
Sampling Strategy for Target Tissue Landmarks with Spatial Domain Associations. A: The procedure to obtain real spatial transcriptomics data from mice with difference months of ages (4 weeks, 24 weeks, 20 months) by the spatial omics technology, MERFISH. B: The simulation procedure for TTLs: generating a TTL with probability 20% (p=20%) around each cell in the selected spatial domain D. Simulate the generation of TTLs for all slices. C: An example for cell type and TTL maps of a slice. D: Example of the cell type and the TTL maps in an FOV sampling sequence in color with unsampled part in grey. E: The Sampling frequency heatmap on a slice with w/ SOFisher within 1000 trails with randomly initial FOVs. F: The Sampling frequency heatmap on a slice with w/o SOFisher for performance comparison. G: The variation in the number of hits with the increase of steps for the two compared policies of w/ and w/o SOFisher. The p-values (denoted by P) between the hits of the two policies at steps 5, 10, 15, 20, 25, 30, 35, 40, 45, 50. H: The minimum number of FOVs for obtaining a target number (=10) of hits, and the number of the first hit in a sampling sequence for 1000 trails with randomly initial FOVs. The p-values of the two performance indices between w/ and w/o SOFisher are given. I: Comparison in terms of the number of hits between w/ and w/o SOFisher within 10, 20, 30, 40, 50 steps. J: In the table, the second and third lines show the average of the number of hits over 200 trails within each of 10, 20, 30, 40, 50 steps and over the total 1000 trails, and the last line show the improvements of w/ SOFisher over w/o SOFisher.

### Generalizability

To evaluate the generalizability of SOFisher when the tissue structures of training differs from the testing data, we utilized the MERFISH dataset^27^ from Fig. 3. This dataset contains approximately 380,000 cells from 31 slices of the mouse frontal cortex and striatum region, spanning 3 different aging stages: 4 weeks, 24 weeks, and 20 months. Previous reports have shown that the brain exhibits distinct tissue structures across these ages. In contrast to Fig. 3, where the analyses did not consider partitioning the dataset based on aging stages, we assessed whether a SOFisher policy trained on one aging stage can effectively guide experiment design on a different aging stage.

We simulated TTL-spatial domain associations by generating TTLs around cells within the corpus callosum domain with a probability of 50% (Fig. 3B). We then trained 4 different SOFisher policies using 4 different training datasets: slices from 4 weeks (SOFisher-4w), 24 weeks (SOFisher-24w), 20 months (SOFisher-20m), and all slices combined (SOFisher-all). To assess the performance of these policies, we evaluated them along with the random sampling strategy on slices from each aging stage: 4 weeks (Fig. 4A), 24 weeks (Fig. 4B), and 20 months (Fig. 4C). The results consistently demonstrated that all the SOFisher policies outperformed random sampling, even in cases where the training and testing datasets were inconsistent, e.g., SOFisher-24w testing on 4-week slices, and SOFisher-20m testing on 24-week slices. These findings suggest that SOFisher exhibits generalizability even when there is a domain gap between training and testing data.

**Fig. 4.**
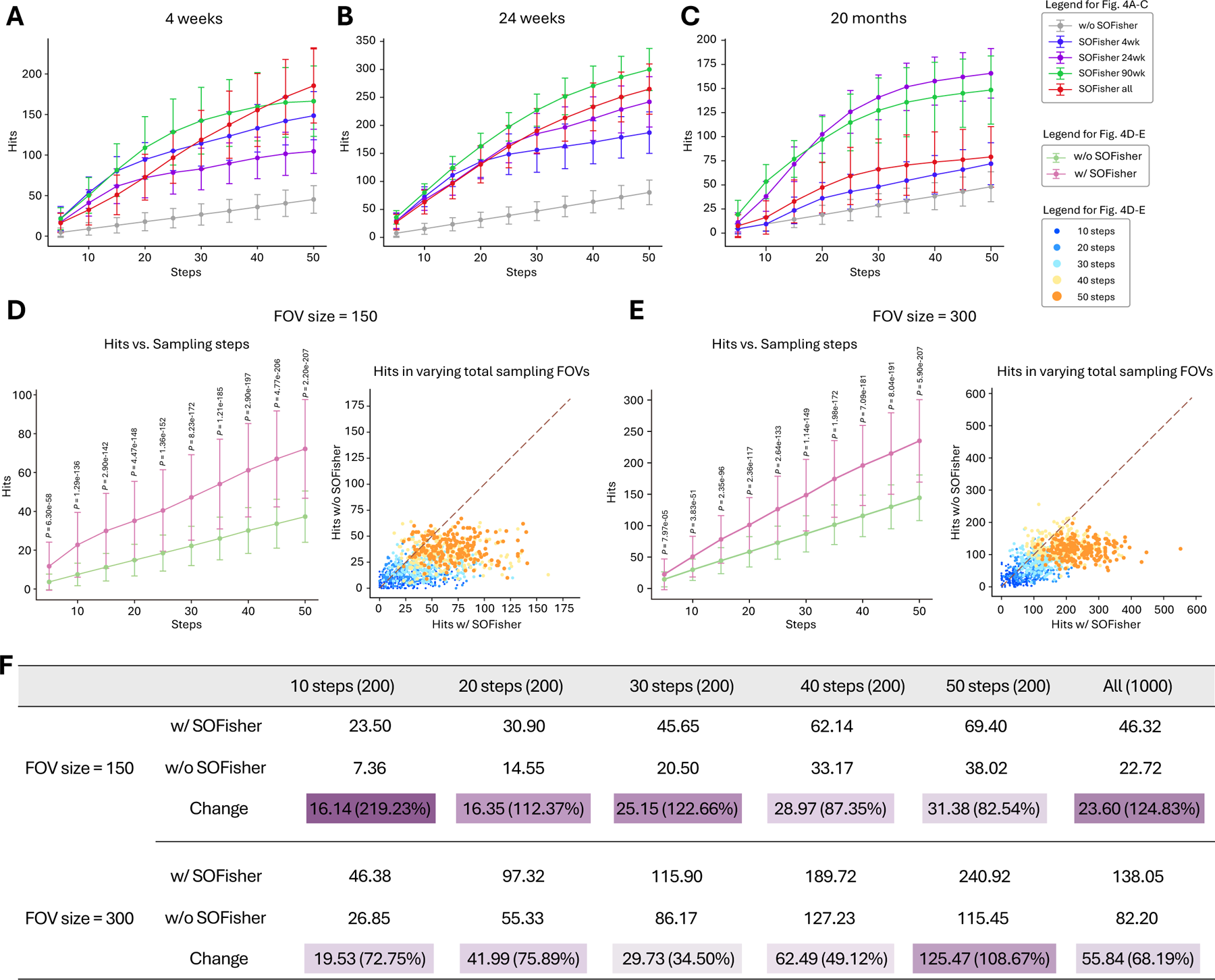
Generalizability and Compatibility. A: The variation in the number of hits with the increase of steps tested on slices from mice with age of 4 weeks, under w/o SOFisher, and SOFisher policy trained by data from mice with ages of 4 week (SOFisher 4wk), 24 week (SOFisher 24wk), 90 week (SOFisher 90wk) and all mice (SOFisher all). B: The variation in the number of hits with the increase of steps tested on slices from mice with age of 24 weeks, under w/o SOFisher, and SOFisher policy trained by data from mice with ages of 4 week (SOFisher 4wk), 24 week (SOFisher 24wk), 90 week (SOFisher 90wk) and all mice (SOFisher all). C: The variation in the number of hits with the increase of steps tested on slices from mice with age of 90 weeks, under w/o SOFisher, and SOFisher policy trained by data from mice with ages of 4 week (SOFisher 4wk), 24 week (SOFisher 24wk), 90 week (SOFisher 90wk) and all mice (SOFisher all). D: For FOVs with the size of 150, the variation in the number of hits with the increase of steps tested on slices from all the mice under w/o SOFisher and SOFisher all, and the comparison in terms of the number of hits between w/o SOFisher and SOFisher all within 10, 20, 30, 40, 50 steps. E: For FOVs with the size of 300, the variation in the number of hits with the increase of steps tested on slices from all the mice under w/o SOFisher and SOFisher all, and the comparison in terms of the number of hits between w/o SOFisher and SOFisher all within 10, 20, 30, 40, 50 steps. F: For FOVs with the size of 150 and 300, the table listing the average of the number of hits over 200 trails within each of 10, 20, 30, 40, 50 steps and over the total 1000 trails, and the last line show the improvements of SOFisher all over w/o SOFisher.

### Compatibility with Different FOV Sizes

In real-world applications, various spatial technologies employ different FOV sizes depending on experimental settings and purposes. To demonstrate SOFisher’s compatibility with different FOV sizes, we tested SOFisher using other FOV sizes such as 150 and 300 (Fig. 4D-F). For each FOV size, SOFisher was applied to the same dataset used in Fig. 2, in which FOV size of 100 was selected. We compared the performance of SOFisher and random sampling by evaluating the number of hits (i.e., the number of TTL-containing FOVs) for a fixed number of FOV sampling steps (10, 20, 30, 40, and 50 steps). The comparison was repeated for 200 times for each number of FOV sampling steps to ensure robustness. Consistent with previous results, SOFisher outperformed random sampling in nearly all cases, regardless of the FOV size and the number of sampling steps (Fig. 4D-F). This finding suggests that SOFisher is compatible with different FOV sizes and can effectively adapt to various real-world experimental settings.

### Sampling Strategy for p-tau in Alzheimer’s Disease

We applied SOFisher to a real dataset of Alzheimer’s Disease (AD) obtained using spatially resolved transcript amplicon readout mapping with protein localization and unlimited sequencing (STARmapPLUS)^28^. This advanced spatial technology measures spatially resolved gene expressions along with two neuropathologic hallmarks^29^ of AD: amyloid-β plaques (Aβ) and neurofibrillary tangles (p-tau) in brain tissues of an AD mouse model at 8 and 13 months of age. Such single-cell multi-omics spatial co-profiling technology remains inaccessible to most labs, and mainstream spatial technologies can only measure either spatially resolved genes or proteins in a tissue at single-cell resolution. Training a SOFisher policy can aid experiment design focused on local niches around Aβ and/or p-tau in spatial single-omics studies.

We ran SOFisher on a single slice from a 13-month-old mouse (Fig. 5A) and visualized a sampling sequence on the cell type and p-tau maps (Fig. 5B). The SOFisher-frequency heatmap, generated from 1000 times, closely matched the p-tau localizations (Fig. 5C) and showed better coverage of p-tau regions compared to random sampling (Fig. 5D). SOFisher consistently outperformed random sampling by achieving a higher number of hits at varying sampling steps (Fig. 5E) and requiring significantly fewer steps to obtain the target number of p-tau-containing FOVs (Fig. 5F). In fixed-step comparisons, SOFisher obtained a larger number of p-tau-containing FOVs (Fig. 5G, H) across varying total sampling steps.

**Fig. 5.**
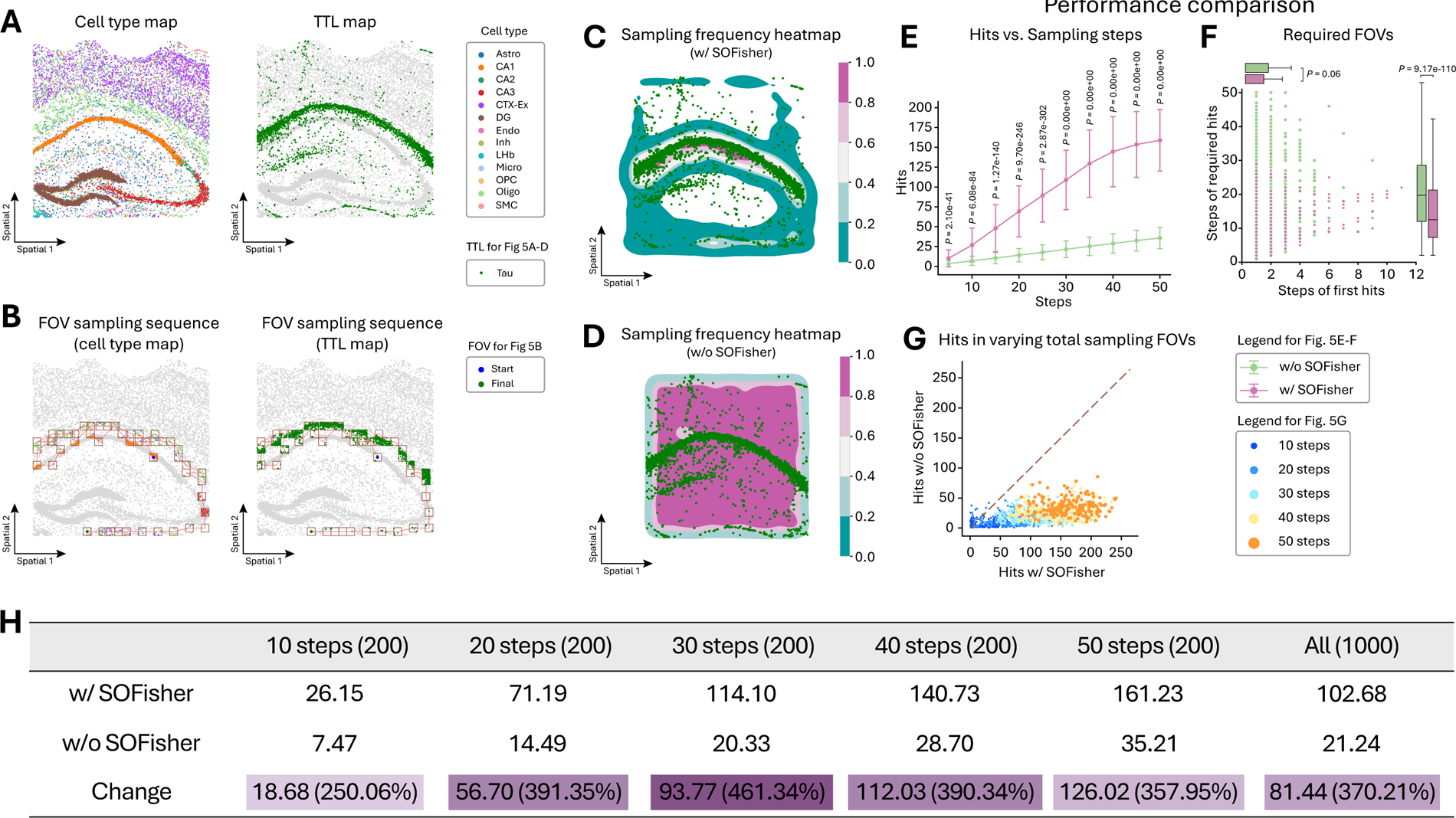
Sampling Strategy for P-tau in Alzheimer’s Disease. A: An example for real data of cell type map and TTL map. B: Example of the cell type and the TTL maps in an FOV sampling sequence in color with unsampled part in grey. C: The Sampling frequency heatmap on a slice with w/ SOFisher within 1000 trails with randomly initial FOVs. D: The Sampling frequency heatmap on a slice with w/o SOFisher for performance comparison. E: The variation in the number of hits with the increase of steps for the two compared policies of w/ and w/o SOFisher. The p-values (denoted by P) between the hits of the two policies at steps 5, 10, 15, 20, 25, 30, 35, 40, 45, 50. F: The minimum number of FOVs for obtaining a target number (=10) of hits, and the number of the first hit in a sampling sequence for 1000 trails with randomly initial FOVs. The p-values of the two performance indices between w/ and w/o SOFisher are given. G: Comparison in terms of the number of hits between w/ and w/o SOFisher within 10, 20, 30, 40, 50 steps. J: In the table, the second and third lines show the average of the number of hits over 200 trails within each of 10, 20, 30, 40, 50 steps and over the total 1000 trails, and the last line show the improvements of w/ SOFisher over w/o SOFisher.

### Sampling Strategy for p-tau and Aβ in Alzheimer’s Disease

To demonstrate SOFisher’s ability to aid experiment design when simultaneously interested in two classes of TTLs, we extended the model to sample FOVs containing both TTLs (see Methods). We designed a third simulation dataset based on the mouse primary motor cortex containing 64 slices^17^ (Fig. 2A). For each slice, we generated two classes of TTLs: TTL1 around LT 45 cells with a 50% probability and TTL2 around LT 23 cells with a 50% probability, simulating TTL1-cell-type and TTL2-cell-type associations, respectively.

The extended SOFisher model successfully guided sampling to FOVs containing both TTL1 and TTL2 (Fig. 6B). The SOFisher-frequency heatmap overlapped well with regions containing both TTL classes (Fig. 6C), outperforming random sampling (Fig. 6D). SOFisher also significantly outperformed random sampling in the number of hits (both-TTL-containing FOVs) at fixed steps (Fig. 6E, G, H), the required steps for the first hit, and the required steps for target hits (Fig. 6F).

**Fig. 6.**
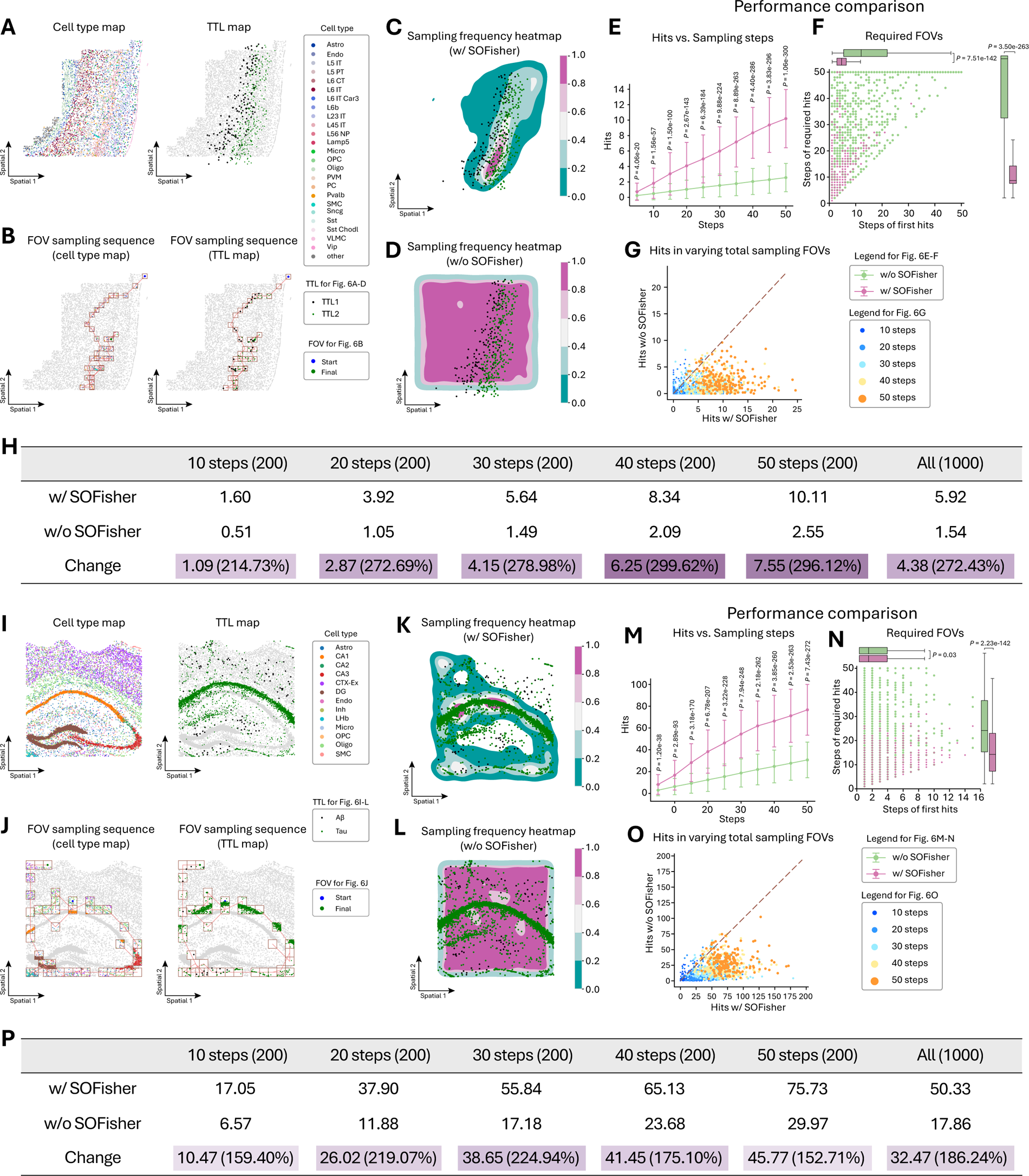
Sampling Strategy for P-tau and Aβ in Alzheimer’s. Disease A: An example for cell type map and two-TTL maps of a slice, while the two types of TTL are generated around two cell types with a probability. B: Example of the cell type and the two-TTL maps in an FOV sampling sequence in color with unsampled part in grey. C: The Sampling frequency heatmap on a slice with w/ SOFisher within 1000 trails with randomly initial FOVs. D: The Sampling frequency heatmap on a slice with w/o SOFisher for performance comparison. E: The variation in the number of hits for both types of TTLs with the increase of steps for the two compared policies of w/ and w/o SOFisher. The p-values (denoted by P) between the hits of the two policies at steps 5, 10, 15, 20, 25, 30, 35, 40, 45, 50. F: The minimum number of FOVs for obtaining a target number (=10) of hits for both types of TTLs, and the number of the first hit in a sampling sequence for 1000 trails with randomly initial FOVs. The p-values of the two performance indices between w/ and w/o SOFisher are given. G: Comparison in terms of the number of hits for both types of TTLs between w/ and w/o SOFisher within 10, 20, 30, 40, 50 steps. H: In the table, the second and third lines show the average of the number of hits over 200 trails within each of 10, 20, 30, 40, 50 steps and over the total 1000 trails, and the last line show the improvements of w/ SOFisher over w/o SOFisher. I: An example for real data of cell type map and two-TTL maps with Abeta and tau. J: Example of the cell type and the two-TTL maps in an FOV sampling sequence in color with unsampled part in grey. K: The Sampling frequency heatmap on a slice with w/ SOFisher within 1000 trails with randomly initial FOVs. L: The Sampling frequency heatmap on a slice with w/o SOFisher for performance comparison. M: The variation in the number of hits for both types of TTLs with the increase of steps for the two compared policies of w/ and w/o SOFisher. The p-values (denoted by P) between the hits of the two policies at steps 5, 10, 15, 20, 25, 30, 35, 40, 45, 50. N: The minimum number of FOVs for obtaining a target number (=10) of hits for both types of TTLs, and the number of the first hit in a sampling sequence for 1000 trails with randomly initial FOVs. The p-values of the two performance indices between w/ and w/o SOFisher are given. O: Comparison in terms of the number of hits for both types of TTLs between w/ and w/o SOFisher within 10, 20, 30, 40, 50 steps. P: In the table, the second and third lines show the average of the number of hits over 200 trails within each of 10, 20, 30, 40, 50 steps and over the total 1000 trails, and the last line show the improvements of w/ SOFisher over w/o SOFisher.

We then trained a SOFisher policy on the real AD mouse dataset^28^ containing spatially resolved gene expression, Aβ, and p-tau on 8-month (2 slices of AD and 2 slices of WT) and 13-month (2 slices of AD and 2 slices of WT) mice. The policy was trained on AD slices to sample FOVs containing both Aβ and p-tau. We evaluated the policy on a 13-month AD mouse brain slice, visualizing the cell types, Aβ, and p-tau (Fig. 6I). Aβ and p-tau were distributed differently: Aβ was prominent in the cortex and hippocampus, while p-tau was strongest in the CA1 region of the hippocampus, with only small portions close together. The sampling sequence (Fig. 6J) under the SOFisher policy showed that the sampled FOVs were not just densely located in the CA1 region, as in Fig. 5B, C, where only p-tau was targeted. Instead, the sampled FOVs tended to cover regions where both p-tau and Aβ occurred (Fig. 6K), outperforming random sampling (Fig. 6L). Quantitative results demonstrated SOFisher’s significantly better performance in the number of hits (Aβ-p-tau-containing FOVs) at fixed steps (Fig. 6M, O, P), the required steps for the first hit, and the required steps for the target hit (Fig. 6N).

### SOFisher-designed experiments capture biological insights in Alzheimer’s Disease

In studying Alzheimer’s Disease (AD) mechanisms and progression, scientists often focus on the local cell niche surrounding Aβ and p-tau^29^. We demonstrate that the trained SOFisher policy, as described in the previous section, can design experiments that reveal broader biological insights, including AD-associated differential abundance of cell types, subtypes, and gene programs. Our aim is to show that SOFisher-designed experiments can achieve comparable or even superior power using small-area, spatial single-omics data compared to large-area, spatial dual-omics experiments. To this end, we conducted two experimental settings for comparative analysis of AD vs. WT mouse tissue. For “Original design”, we employed entire AD slices as the condition dataset and entire WT slices as the control dataset, with spatial dual-omics STARmapPLUS (spatial gene expression and protein co-profiling). For “SOFisher design”, we used SOFisher-sampled FOVs as the condition dataset and entire WT slices as the control dataset, with only spatial gene expression data (a common scenario in real-world applications).

We first performed differential abundance (DA) analysis using data from Original design and SOFisher design, respectively. We selected MELD^30^ for DA analysis based on a recent benchmarking study^31^. MELD was used to estimate the relative likelihood of each cell between AD and WT conditions (see Methods). The SOFisher design (Fig. 7B, D) highlighted AD-associated cell types with relative likelihood > 0.5 consistent with the original design (Fig. 7A, C), including astrocytes (Astro), endothelial cells (Endo), microglia (Micro), oligodendrocyte precursor cells (OPC), oligodendrocytes (Oligo), and smooth muscle cells (SMC). Intriguingly, in the SOFisher design, we observed a wide range of AD relative likelihood values in Astro, Micro, and Oligo (Fig. 7D), suggesting the presence of subtypes with different perturbation responses to AD, consistent with previous studies^32–35^. This wide range is more evident in the SOFisher design DA analysis (Fig. 7D) than in the original design (Fig. 7C).

**Fig. 7.**
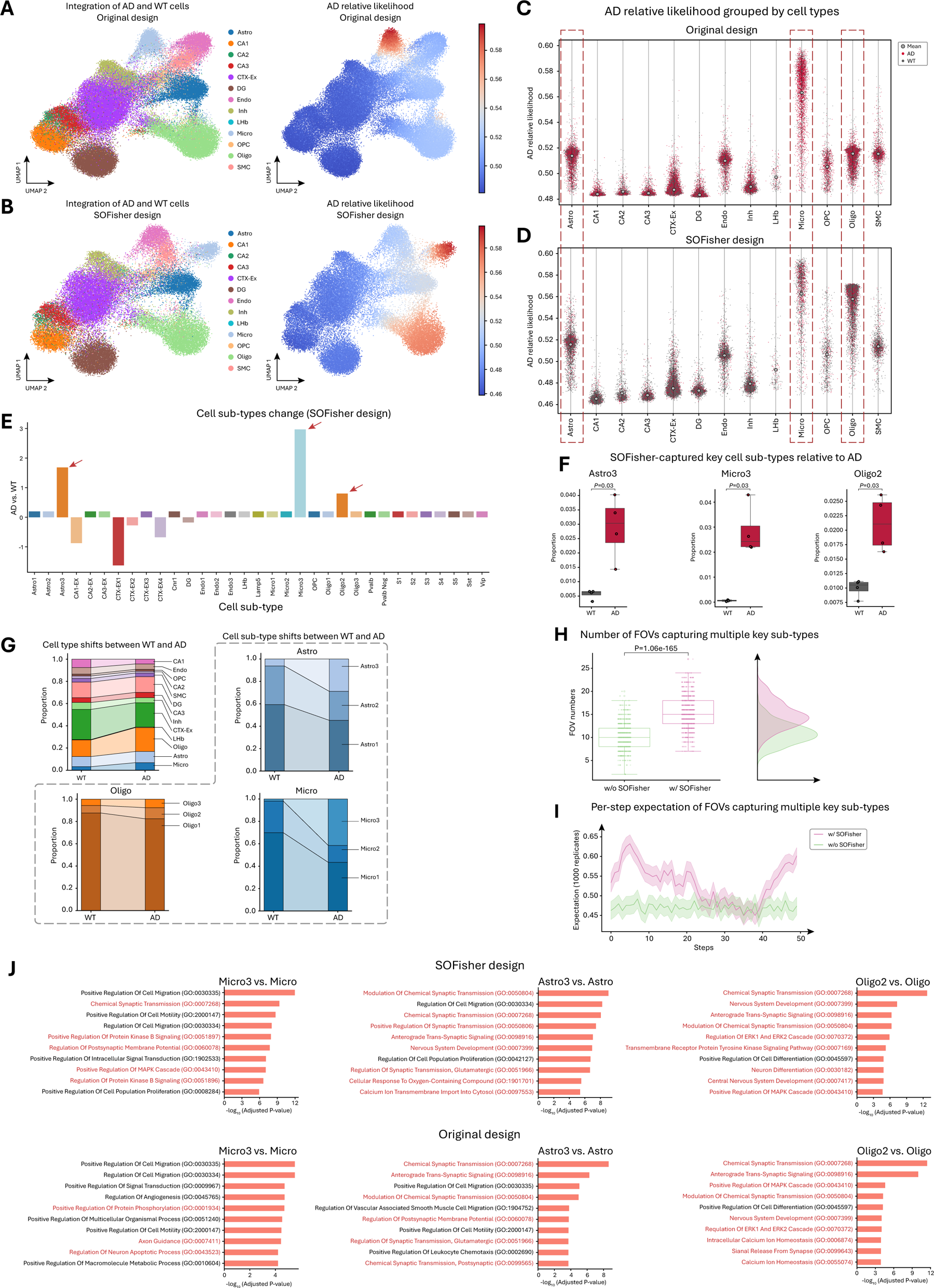
Biological insight discovery with SOFisher at cell and gene levels. A: UMAP visualization of the original design (entire AD versus entire WT) embedding after batch integration across different replicates in AD and WT conditions. Colors indicate cell type and AD-relative likelihood estimated by MELD for each cell, with higher likelihood indicating greater AD relevance. B: UMAP visualization of SOFisher design (SOFisher sampled FOV versus entire WT) embedding after batch integration across different replicates in AD and WT conditions. C: Visualization of AD relative likelihood for each cell in the original design, grouped by cell types across AD and WT. Scatter points indicate mean relative likelihood values in each cell group, with higher relative likelihood indicating greater AD relevance. Original design shows high relative likelihood mean value only in Microglia. D: Visualization of AD relative likelihood for each cell in SOFisher design, grouped by cell types across AD and WT. Scatter points indicate mean relative likelihood values in each cell group. SOFisher exhibits both high relative likelihood mean value and dispersion among three key cell types (Astrocyte, Microglia, Oligodendrocyte), outperforming the original design. E: Visualization of cell subtype proportion changes. Proportion change is defined by the log2 fold change ratio estimated by scCODA under 0.2 FDR control. Higher change ratio indicates higher proportion in AD groups. Statistical significance is generated considering each of the eight slices as replicates during scCODA estimation. Key cell subtypes (Micro3, Oligo2, Astro3) exhibit higher proportions in AD groups. F: SOFisher-captured key cell subtypes (Micro3, Oligo2, Astro3) relative to AD across eight replicates (four in AD and four in WT). Proportion in AD groups is calculated on cells in SOFisher sampled FOVs for each slice, while WT cell proportion is calculated in the entire slice. SOFisher design shows significantly higher proportion in all three key cell subtypes (P < 0.05, rank-sum test). G: Cell type and subtype proportion change tendencies in the SOFisher-designed experiment across all samples. Left-upper: main cell type proportion change from WT to AD; remaining three: cell subtype proportion change for each cell type. The three key cell subtypes (Micro3, Astro3, Oligo2) show the largest ratio increase in the three main cell types. H: Number of FOVs capturing multiple key sub-types in each experiment (50 steps/FOVs) with 1000 random replicate experiments. SOFisher strategy shows significantly higher numbers than baseline (P < 0.05, rank-sum test). I: Per-step expectation of FOVs capturing multiple key sub-types. Comparison of SOFisher design strategy with baseline random strategy shows SOFisher design is non-trivial and biologically meaningful. Line chart shows results of 1000 random replicated experiments with mean and 95% confidence interval; SOFisher design maintains a stable and high capture ratio R (see Methods, Sampling strategy comparison) compared to baseline. J: GO term enrichment analysis results on original design and SOFisher design for significant differential gene sets (FDR adjusted P < 0.05, log2 fold change ratio > 1) for key cell subtypes in AD versus main cell types in WT in each experiment design. AD-related terms are shown in red.

For a more fine-grained analysis considering slice replication with statistical significance, we employed the Bayesian inference-based scCODA^36^ to determine significantly enriched cell subtypes in AD using the SOFisher design experiment (see Methods). We investigated whether the SOFisher design could capture cell subtypes as reported in the original design. Fig. 7E shows that the SOFisher design exhibited significantly higher abundance in three cell subtypes (‘Astro3’, ‘Micro3’, ‘Oligo2’) compared to WT samples. These subtypes were previously recognized as key populations closely associated with AD. Furthermore, we demonstrated that these identified cell subtypes exhibited significant abundance differences between WT and AD using only SOFisher design data, both within each slice (Fig. 7F) and collectively (Fig. 7G).

We further demonstrated that the SOFisher design can better capture the three key cell subtypes in AD (i.e., ‘Astro3’, ‘Micro3’, ‘Oligo2’). We compared the number of FOVs capturing all three key cell subtypes in 50 FOV-sampling steps with and without the aid of SOFisher. A higher number of FOVs capturing all three key cell subtypes were obtained with SOFisher than without it (Fig. 7H). The per-step expectation (1000 replicates per step) of capturing all 3 key cell subtypes also showed substantial improvement with SOFisher (Fig. 7I). This indicated that SOFisher can precisely locate the cellular phenotypes surrounding Aβ and p-tau even when researchers have only a limited vision of the whole tissue, and the visible information (i.e., cellular phenotypes within FOVs) does not contain any Aβ or p-tau information.

Finally, we demonstrated that the SOFisher-captured key subtypes can reveal more AD-related gene ontology (GO) terms (adjusted p-value < 0.05, log2 fold change ratio > 1) than the original design. For the SOFisher design, we performed GO enrichment analysis using differentially expressed genes obtained from comparing SOFisher-captured Micro3 and WT Micro (Fig. 7J top). We also performed similar analyses on Astro and Oligo (Fig. 7J top). For the original design, we performed GO enrichment analysis using differentially expressed genes obtained from comparing all Micro3 in AD and WT Micro (Fig. 7J bottom). We also performed similar analyses on Astro and Oligo (Fig. 7J bottom). We highlighted GO terms closely related to AD biological processes in red, and the SOFisher design contained more related GO terms than the original design.

## Discussion

Spatial Omics is rapidly advancing in both experimental^5,6,9,19,37–43^ and computational^44–50^ domains. We introduce SOFisher, a reinforcement learning-based framework designed to optimize field of view (FOV) sampling strategies in spatial omics experiment designs. The results demonstrate that SOFisher significantly improves the efficiency of capturing regions of interest (ROIs) compared to conventional sampling approaches. The performance of SOFisher across various simulated datasets and its successful application to a real Alzheimer’s Disease (AD) dataset underscore its potential to revolutionize spatial omics experiment design. By leveraging information from previously sampled FOVs, SOFisher effectively guides the selection of subsequent FOV positions, leading to more targeted and efficient data collection. This approach not only saves time and resources but also enhances the quality of data obtained from spatial omics experiments.

One of the key strengths of SOFisher is its ability to adapt to different experimental settings. The framework’s generalizability, demonstrated through its performance across different aging stages of mouse brain datasets, suggests that SOFisher can be applied to cases with domain gaps. The compatibility of SOFisher with different FOV sizes further emphasizes its practicality in real-world experimental settings. This flexibility allows researchers to tailor their sampling strategies to specific technical constraints or biological questions.

The application of SOFisher to the AD dataset yields particularly promising results. By guiding the selection of FOVs containing both AD pathology markers, SOFisher enables the identification of key differential abundance cells, subtypes, and gene programs related to AD pathology. This achievement is significant, as it demonstrates that SOFisher can facilitate discoveries typically associated with spatial multi-omics experiments on large areas, using only a spatial single-omics experiment on a smaller area. This capability has the potential to democratize access to advanced spatial biology insights, making them achievable with more widely available and cost-effective technologies.

SOFisher addresses three critical gaps in current spatial omics experiment designs. Firstly, it mitigates the high cost associated with whole tissue coverage by optimizing FOV selection. Secondly, it overcomes the challenge of FOV selection with limited prior knowledge of the tissue slice to be measured. Lastly, it provides a practical alternative to single-cell spatial multi-omics technologies, which are often infeasible due to technical or resource constraints.

Another important feature of SOFisher that warrants further discussion is its flexibility, particularly in utilizing different types of cellular phenotype information. Although in this study SOFisher observed cell type as the primary cellular phenotype, it can be easily adjusted to leverage other cellular characteristics, such as gene expression patterns, to infer the next position of FOV.

Despite these promising results, several limitations and areas for future research should be acknowledged. Due to limited available spatial multi-omics data at single-cell resolution, the performance of SOFisher on the complexity and heterogeneity of the tissue has not been comprehensively studied. Future work should focus on evaluating and optimizing SOFisher’s performance across a broader range of tissue types and pathological conditions. Additionally, the integration of SOFisher with other experiment design considerations, such as target selection and multiplexing strategies, could further enhance its utility in spatial omics research.

We can also vision future extensions of SOFisher in terms how to award the target FOVs. For example, SOFisher could be adapted to target FOVs that are most predictable by H&E staining (or other cheap assays)^51,52^, enabling researchers to predict gene expression patterns across an entire tissue slice with H&E using only a limited number of sampled FOVs as training data. Similarly, the framework could be extended to sample FOVs containing highly informative gene expression data, allowing researchers to selectively sample key areas using expensive but comprehensive assays (e.g., covering large portion or whole transcriptome) and then impute whole transcriptome profiles for unmeasured regions^53,54^. Another important avenue for future research is the exploration of SOFisher’s potential in longitudinal studies^55^. Adapting the framework to guide sequential sampling over time could provide valuable insights into dynamic biological processes and disease progression.

## Methods

### Reinforcement Learning (RL) based experiment design

Under the general framework of RL, an agent in a state *s_k_* selects an action *a_k_* at each time step *k*. Then, the environment responds to that state-action pair with a reward *r_k_*, and presents a new state *s_k+1_* to the agent. This sequential decision-making scenario is formalized as a Markov decision process (MDP), denoted by the tuple 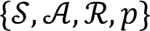, where & is the set of states (state space), S is the set of actions (action space), ℛ is the reward space, and 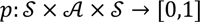 is the transition probability distribution.

The agent aims to learn an optimal policy π^∗^ by maximizing an expected cumulative reward, i.e.,

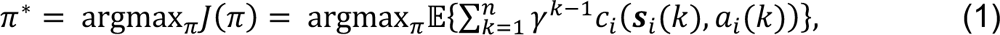

where 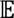 denotes the expectation operator, 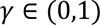 denotes a discounting factor, 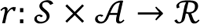 is a reward function, and 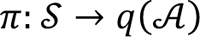 is a policy function mapping a state to a probability distribution over the actions. There are two main categories of methods to learn an optimal policy in (1): i) *value function* methods and ii) *policy gradient* methods. In this work, a TTL searching algorithm based on value function methods is designed in the following sections.

### MDP modeling

Before the MDP modeling, we divide each tissue slice into an *m_r_* x *m_c_* grid, where the size of each grid is predefined. Each FOV is assumed to be a square with a side length of *r_s,_* its sampling center is located at a normalized position *p* = [*x*, *y*] based on the body axis system with the origin at the center of the slice. Thus, a searching policy only aims to find a sequence of sampling positions such that the largest number of TTL can be searched by the FOVs.

The formalization of a finite MDP requires the definition of 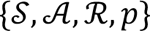. When solving an MDP with a model-free RL method, the last element p is considered unknown, and the three remaining elements 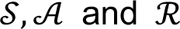 are defined as follows.

### The state

We select the state *s_k_* containing three parts: the FOV’s sampling position *p_k_* at the current time step *k*, a *n*-dimensional row vector with each element as the number of a type of cell in the FOV, i.e., 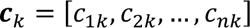, and a row vector *m_k_* representing the search grade in the slice. That is,

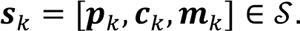

We elaborate the grade vector *m_k_* by leveraging an adaptive partition method. To avoid repeated search, we indicate each grid cell with 1 as searched and 0, otherwise, such that a matrix with *m_r_ x m_c_* elements can be obtained to represent the search grade of the whole slice. Nonetheless, the challenge is that the number of elements vary with the size of a slice. To overcome it, we propose the adaptive partition method, and define the grade vector *m_k_* with 17 dimensions. The first 9 elements are the indicated values of the grid cells:

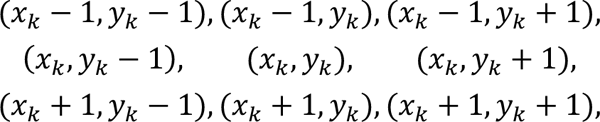

and the last 8 elements are average indicated values within each of 8 domains

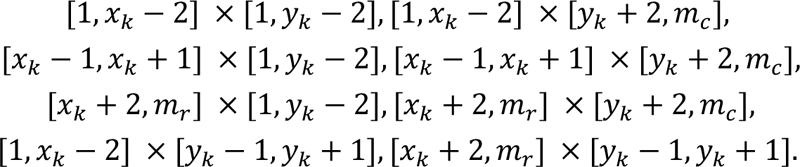

### The action

For the TTL searching problem, we set the action as a single value 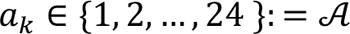. By executing an action, the FOV moves to one of the 24 sampling positions closed to its current 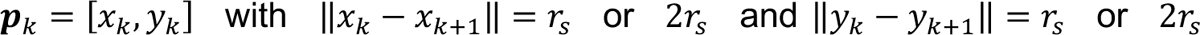 in Supplementary Fig. 3, where each sampling position corresponds to an element of 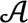. Clearly, any two adjacent samples do not overlap except the cases of the FOV encountering the boundaries of the slice.

In practice, the sampling center of the next FOV can be selected at any position over the slice, which implies an *m_r_* x *m_c_*-dimensional action space. In comparison, we intuitively design the next FOV closed to the current one due to the fact that the distribution of TTL is relatively continuous. In another word, if we can detect TTL at position *p_k_*, then it is likely to detect more around *p_k_*. In addition, our 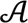 with 24 dimensions can greatly reduce the difficulty in learning an optimal policy.

### The reward

Considering the objective of maximizing the number of TTLs, we need to design a reward function to guide the policy learning for informative sample sequence. To this end, the reward function at time step *k* is defined as

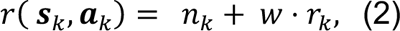

where *n_k_* is the number of TTLs in the current FOV and denotes the reward for searching TTLs; *r_k_* is the number of times to resample the current grid cell and thus its weighting coefficient *w* is a negative constant representing a penalty. With all these definitions in place, a reward can be easily computed after executing action *a_k_* with state *s_k_*.

### Double DQN-based RL Algorithm

To solve the MDP in the last section, this section proposes the TTL search algorithm based on double deep Q-network (DQN)^56^.

We give a quick glance of the classical DQN^57^, which utilizes a deep neural network *Q_n_*. parameterized by a set of weights η to approximate the state-action value function 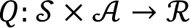, i.e., Q-function. The optimal Q-function is formally defined as

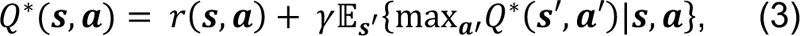

Where (*s*’, *a*’) denotes the state and action taken after taking action *a* in state *s.* DQN aims to converge to the optimal Q-function in (3), and to find the associated optimal stationary policy as

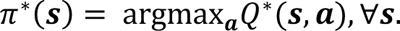

Moreover, the weights η are updated with the gradient of the loss function

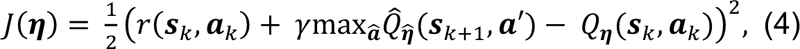

where the tuple 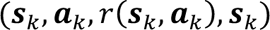 is sampled from a replay memory *D* that stores historical data, and 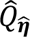 is a target Q-network parameterized by weights 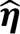. DQNs can stabilize the learning algorithm thanks to the techniques of experience replay and target Q-network^58^.

Compared with the classical DQN, the double DQN replaces the greedy term 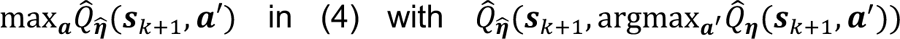, which mitigates the overestimation of the values of the Q-function associated with classical DQN designs. Thus, at each iteration, the weights of the Q-network are updated with the gradient of the loss function

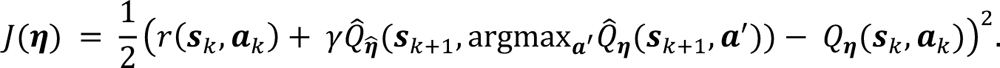

With all this in mind, an RL algorithm is proposed to address the TTL search problem based on the double DQN, which is summarized in Algorithm 1. In essence, the algorithm blends the steps of a double DQN with the observation and tracking of the states and measurements.

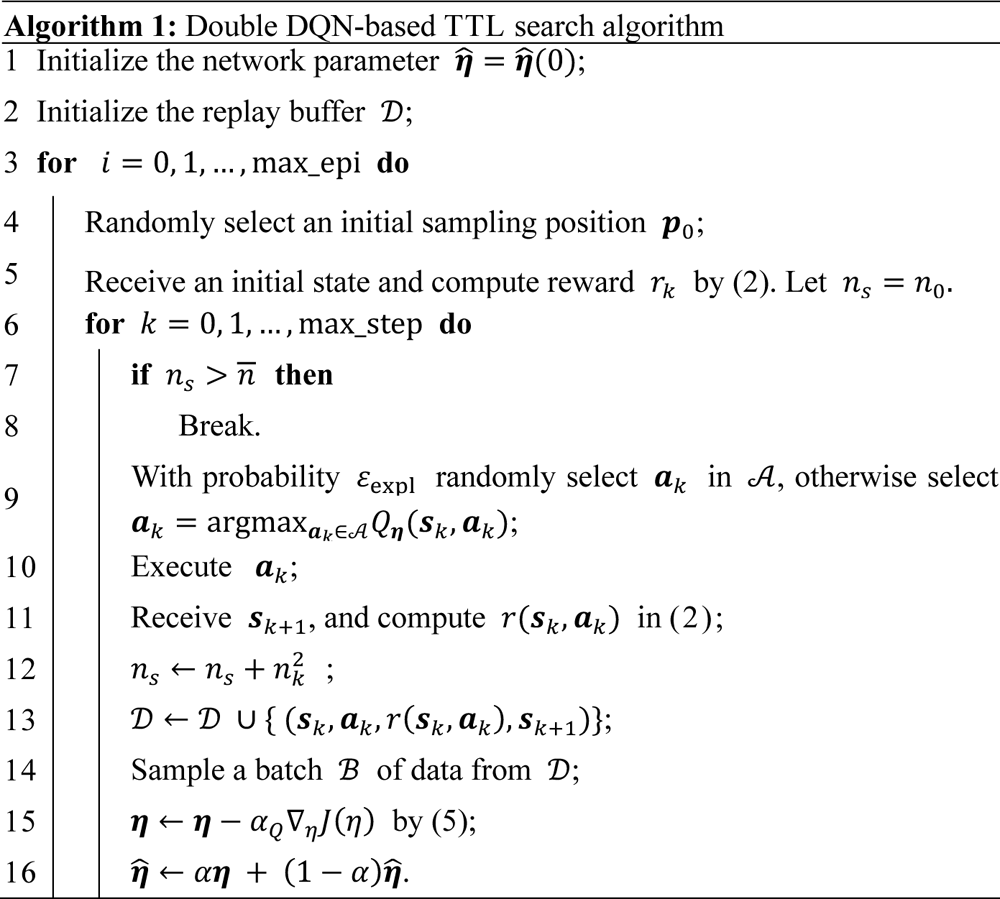

The algorithm involves a parameter 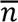 enoting a terminal condition of a training episode. It implies that sampling can stop if sufficient TTL has been found. This setting can boost the convergence of the policy learning. Moreover, we consider max_epi number of episodes and max_step time steps per episode. At each time step *k*, we store 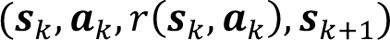 in a replay buffer *D* (cf. line 13). The network is trained by a batch ℬ of data randomly sampled from D (cf. line 14). A greedy policy is adopted with exploration probability $\epsilon$, which decreases with the iterations at a linear rate with an exploration parameter decay. The stochastic gradient descent technique Adam^59^ is used to minimize (5) with learning rate *α_Q_*. Moreover, a soft update is employed in line 16 for the weights 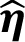 of the target network, using a weight *α*. As already mentioned, line 7 checks if the conditions based on 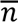 is satisfied and, if yes, the episode is terminated.

## Experiments

### Environment Details

We have trained the TTL search policies by three datasets, namely, Cortex (used in Fig. 2), Aging (used in Fig. 3, 4), and Control (used in Fig. 5, 6). While the first two datasets only contain distributions of various cells, the last dataset contain distributions of cells and TTL proteins. Thus, we simulate the TTL distributions in Cortex and Aging according to some features. To be specific, for Cortex, TTL is generated closed to certain types of cells with probabilities; and for Aging, TTL is generated associated within certain anatomical region with probabilities. The detailed selections are given in Table I.

**TABLE I.**
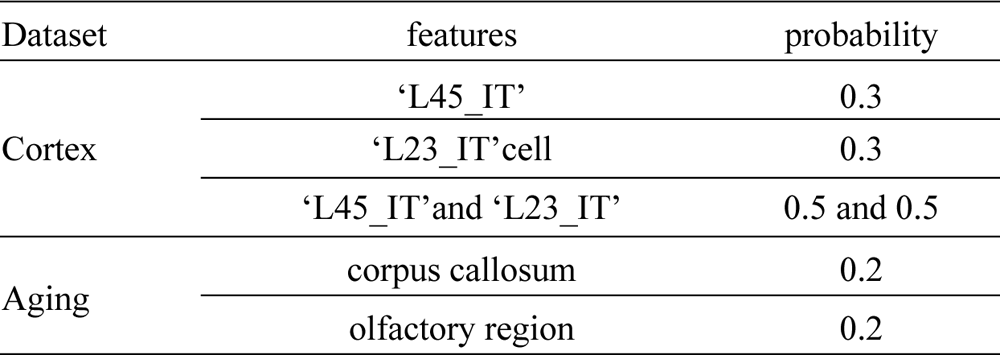
Detains for generation.

For the state space S of the MDP, we used cell types as cellular phenotypes for the three datasets in Table II.

**TABLE II.** Cell types for the state space.

| Dataset | Cell types |
| --- | --- |
| Cortex<br>(21 cells) | ‘L45_IT’, ‘L23_IT’, ‘L56_NT’, ‘L5_IP’, ‘L5_PT’, ‘Lamp5’,<br>‘L6_CT’, ‘L5_IT’, ‘L6_IT’, ‘L6JT_Car3’, ‘L6b’, ‘OPC’,<br>‘Astrocytes’, ‘Endothelial’, ‘Oligodendrocytes’, ‘Microglia’,<br>‘VLMC’, ‘Sst’, ‘SMC’, ‘Pvalb’, ‘Pericytes’, ‘PVM’ |
| Aging<br>(14 cells) | ‘Astro’, ‘Endo’, ‘Epen’, ‘ExN’, ‘InN’, ‘MSN’, ‘OPC’,<br>‘Micro’, ‘Macro’, ‘Olig’, ‘Peri’, ‘T cell’, ‘Vlmc’ |
| AD<br>(13 cells) | ‘Astro’, ‘Micro’, ‘CTX-Ex’, ‘DG’, ‘Endo’, ‘Oligo’,<br>‘CA1’, ‘CA2’, ‘CA3’, ‘Inh’, ‘SMC’, ‘LHb’, ‘OPC’ |

For the reward function in (2), we set the weighting parameter as 5. In particular, in the case of jointly searching two types of proteins, the reward term *n_k_* is defined as the product of the number of Aβ the number of p-tau.

In addition, the parameters in Algorithm 1 vary with the datasets, and we list them in the following Table III.

**TABLE III.**
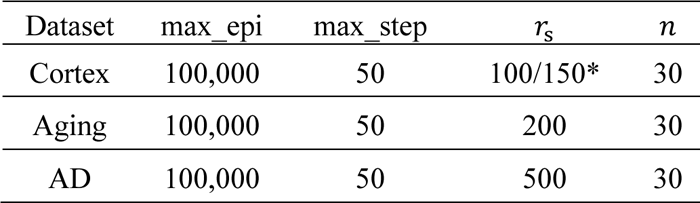
Parameters in Algorithm 1.

| Dataset | max_epi | max_step | $r_s$ | $n$ |
| --- | --- | --- | --- | --- |
| Cortex | 100,000 | 50 | 100/150* | 30 |
| Aging | 100,000 | 50 | 200 | 30 |
| AD | 100,000 | 50 | 500 | 30 |

See Table IV for the hyper-parameters of Algorithm 1.

**TABLE IV.** Hyperparameters.

| Hyperparameters | Value |
| --- | --- |
| Minibatch size | 256 |
| Learning rate | 0.001 |
| Target smoothing coefficient ($\tau$) | 0.005 |
| Discount ($\gamma$) | 0.99 |
| Exploration parameter ($\epsilon$) | 1 |
| Final exploration parameter ($\epsilon$) | 0.01 |
| Exploration parameter decay | 0.0005 |
| Number of hidden layers | 2 |
| Number of neurons in each hidden layer | 256 |

### Testing Details

We test the double DQN policy learned by Algorithm 2 as follows, where the parameters are similarly selected to those in Table V.

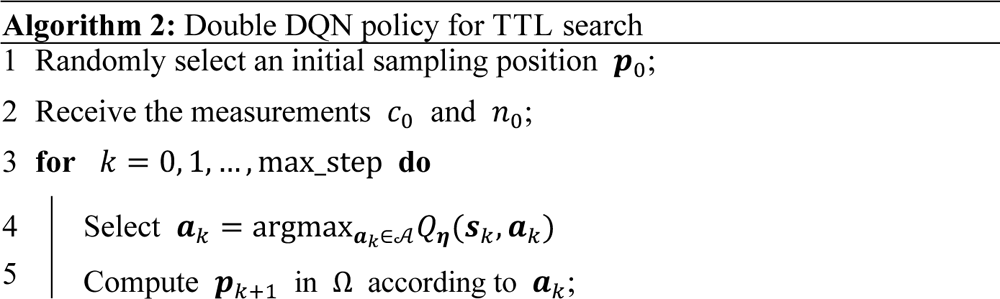
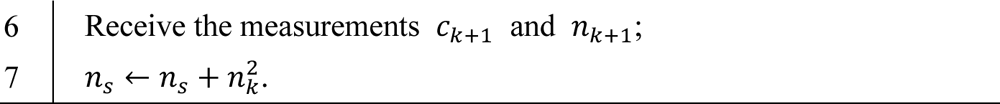

In addition, we select a random policy for TTL search to compare with the Double DQN policy. By the random policy, the sampling position for the FOV is selected within the entire slice range, denoted by Ω, with equal probability at each step. That is, only lines 5-6 in Algorithm 2 need to be replaced by randomly selecting a sampling position 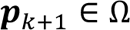. In practice, this policy can be easily implemented. The training and testing data settings are listed in Supplementary Table 1.

### Downstream analysis

Given the SOFisher-designed FOV in a real AD mouse dataset, we conducted a comparative analysis of the downstream outcomes for two condition group settings: (1) all AD slices versus normal slices, and (2) SOFisher-designed FOV on AD slices versus normal slices. This comparison focused on cell type differential abundance and gene expression differential expression to validate that the SOFisher-designed FOV can achieve comparable or superior results compared to the all AD condition splitting setting.

### Differential abundance

We conducted differential abundance analysis at various levels to validate the utilization of SOFisher-designed FOV. Differential abundance analysis at the cell type level reveals changes in cell composition between control and condition groups, identifying key cell types related to disease mechanisms, which are crucial for downstream analysis in real biological experiments.

The first analysis employed is the graph-based condition likelihood estimation (MELD). Given input data comprising both condition and control cells, MELD utilizes a graph diffusion procedure based on cell similarity to estimate the likelihood of each cell belonging to the condition category. For each condition group splitting setting, we conducted the procedure independently as follows: We first utilized Harmony^60^ to integrate the samples into a batch-integrated embedding space, designating AD cells as the condition group and normal cells as the control group. We then applied the MELD estimation procedure to these input cells based on the similarity graph constructed based on the batch-integrated embedding. Higher MELD estimated likelihood indicates greater relevance to the AD group.

However, the MELD likelihood does not provide direct statistical significance considering the replication in the input samples. Therefore, with the key cell types identified by MELD, we conducted scCODA analysis for the SOFisher-designed condition splitting to achieve a finer-grained cell subtype differential abundance analysis with statistical significance. scCODA introduces a Bayesian method to address the common issue of low replicate numbers in single-cell analysis. It employs a hierarchical Dirichlet-Multinomial model to account for uncertainty in cell-type proportions and negative correlation bias by jointly modeling all measured cell-type proportions. Each slice was regarded as a replicate, including both AD and control slices, and only the SOFisher-designed FOV cells in each AD slice were used. We utilized the annotated fine-grained cell subtype as cell type input for scCODA, and the outcome of scCODA is the log fold-change ratio for each cell subtype with False Discovery Rate (FDR) control. After scCODA training, we selected the significantly shifted cell subtypes under an FDR control threshold of 0.2.

### Differential expression and GO term enrichment analysis

Following the cell type-level differential abundance analysis, we applied another critical downstream task in the analysis of condition versus control groups: differential gene expression analysis. Differential expression analysis aids researchers in identifying changes in gene expression patterns between condition and control groups, with significant differentially expressed genes serving as inputs for more granular analyses, such as biological network and gene set enrichment. In our study, we aimed to demonstrate that the SOFisher-designed FOV condition groups can capture AD-related differential gene expression patterns comparable to those observed in the all-slice condition group setting.

For both condition group settings, we conducted the analysis independently using the following procedure: First, we applied normalization and log transformation to obtain normalized gene expression values. Next, we used Combat^61^ in scanpy^62^ to remove batch effects, yielding batch-corrected gene expression values. We then utilized the **scanpy.tl.rank_genes_groups** function with the Wilcoxon test parameter to analyze batch-corrected and normalized gene expression for the three key cell subtypes in condition groups and their corresponding coarse-grained cell types in control groups. Specifically, for each key cell subtype (Astro3, Micro3, Oligo2), we performed differential gene expression analysis between the cell subtype in condition groups and its corresponding coarse-grained cell type (Astrocyte, Microglia, Oligodendrocyte) in control groups. Genes with an adjusted p-value < 0.05 and a log2 fold change > 1 were considered significant. This differential gene expression analysis provided a significant gene set for each cell type in the three key cell subtypes for both condition splitting settings, which served as the input for subsequent GO term enrichment analysis.

We conducted the GO term enrichment analysis using the gseapy package with the enrichr API. The target gene set was GO Biological Process 2023. We considered GO terms with an FDR-adjusted p-value < 0.05 as significantly enriched for both condition splitting results’ significant gene sets.

### Sampling strategy comparison

We compare the SOFisher-designed FOV with randomly chosen FOVs on the same slice, utilizing random replication to demonstrate that the SOFisher outcome is non-trivial and significantly better than a random strategy.

Given one AD slice, we varied the random seed for a trained SOFisher model during the inference period to obtain random replications of the SOFisher FOV results. We also employed a uniform random sampling strategy, using the same number of FOVs per replication and the same number of replications, to generate the baseline random FOVs. We conducted 1000 replications for both strategies, with each replication sampling 50 FOVs (50 steps). The final outcome for both strategies was an array of shape [1000, 50, 2], representing 1000 random sampling strategy replications, each consisting of 50 steps, and each step containing the central point coordinates for the rectangular FOVs on the slices.

Given these replication results, we examined each strategy’s capability to capture the three key cell subtypes (Astro3, Micro3, Oligo2). We used the key cell subtype capture ratio for each step as a measure of the strategy’s biological relevance and convergence. Formally, given the central point coordinate for one step (FOV), we calculated the key cell subtype ratio using a predefined rule. We used *I_A_, I_M_, I_O_* as indicators for whether the FOVs contains the corresponding cell subtype (Astro3, Micro3, Oligo2), respectively. Thus, we can define the key cell subtype capture ratio R = (*I_A_, I_M_, I_O_*) / 3, and each step, we obtained a ratio p. We calculated p for each step in all replications for both strategies and then compared them.

Furthermore, we want to compare the two strategies from a more global perspective. For each replication, we calculate the number of FOVs that captured all three key cell subtypes (p =1) and then obtain two distributions with 1000 samples each for both the SOFisher-designed strategy and the random sampling strategy in terms of the fully captured FOV numbers. We then conduct a hypothesis test to determine whether the number of fully captured FOVs in the SOFisher strategy is significantly larger than that in the random sampling strategy.

### Boxplot

The lower and upper hinges show the first and third quartiles (the 25th and 75th percentiles); the center lines correspond to the median. Boxplots were generated using Seaborn [https://seaborn.pydata.org/].

### Computational resource

All experiments are performed on a PC running Windows 11 with a 13th Gen Intel(R) Core(TM) i9-13900H at 2.60 GHz with RAM 32.00GB. Our implementation employs PyTorch with Python 3.9.0. and torch 1.12.0+CPU.

## Data availability

The MERFISH mouse primary motor cortex data is available at Brain Image Library: [https://doi.brainimagelibrary.org/ https://doi.org/10.35077/g.21], also can be found in SODB^63^ [https://gene.ai.tencent.com/SpatialOmics/dataset?datasetID=30], and can be loaded using Pysodb^64^ [sodb.load_dataset(’zhang2021spatially’)] . The MERFISH mouse aging data is available at CELL x GENE repository [https://cellxgene.cziscience.com/collections/31937775-0602-4e52-a799-b6acdd2bac2e], also can be found in SODB

[https://gene.ai.tencent.com/SpatialOmics/dataset?datasetID=184], and can be loaded using Pysodb [sodb.load_dataset(’Allen2022Molecular_aging’)]. The STARmapPLUS mouse AD and WT data are available at [https://doi.org/10.5281/zenodo.7332091].

## Acknowledgments

Z.Y. acknowledges the support by National Nature Science Foundation of China (62303119), Shanghai Science and Technology Development Funds (23YF1403000), Chenguang Program of Shanghai Education Development Foundation and Shanghai Municipal Education Commission (22CGA02), National Key R&D Program of China (2023YFF1204800), Shanghai Science and Technology Commission Program (23JS1410100), Tencent AI Lab Rhino-Bird Focused Research Program (RBFR2023008), Shanghai Municipal Science and Technology Major Project (No.2018SHZDZX01), ZJ Lab, and Shanghai Center for Brain Science and Brain-Inspired Technology, and 111 Project (No.B18015). Z. L. acknowledges the support by National Nature Science Foundation of China (62303054).

## Author contributions

Z.Y. and Z.L. conceived and designed the study. Z.L., W.W., and Y.C. developed the computational methods and performed the analysis. J.S. provided guidance on optimization. Z.Y., and Z.L. wrote the manuscript.

## Inclusion & Ethics

Not relevant.

---

**Supplementary Fig. 1 Sampling Strategy for Target Tissue Landmarks with Cell Types Associations (another example related to Fig. 2)**

This figure is similar to Fig. 2C-J. The difference is that the TTLs are simulated based on another cell type. A: An example for cell type and TTL maps of a slice. In particular, we extend each slice into a rectangle and arbitrarily select one corner of the rectangle as the origin of a Cartesian coordinate system, with the two sides of the angle being the coordinate axes: Spatial 1 and Spatial 2. B: Example of the cell type and the TTL maps in an FOV sampling sequence in color with unsampled part in grey. C: The Sampling frequency heatmap on a slice with SOFisher policy (w/ SOFisher) for method evaluation, where we record the number of times each position on the slice is sampled within 1000 trails with randomly initial FOVs. D: The Sampling frequency heatmap on a slice with random policy (w/o SOFisher) for performance comparison. E: The variation in the number of TTL-containing FOVs (hits) with the increase of the total number of FOVs (steps) for the two compared policies of w/ and w/o SOFisher. The p-values (denoted by P) between the hits of the two policies at steps 5, 10, 15, 20, 25, 30, 35, 40, 45, 50. F: The minimum number of FOVs for obtaining a target number (=10) of hits, and the number of the first hit in a sampling sequence for 1000 trails with randomly initial FOVs. The p-values of the two performance indices between w/ and w/o SOFisher are given. G: Comparison in terms of the number of hits between w/ and w/o SOFisher within 10, 20, 30, 40, 50 steps. H: The second and third lines show the average of the number of hits over 200 trails within each of 10, 20, 30, 40, 50 steps and over the total 1000 trails, and the last line show the improvements of w/ SOFisher over w/o SOFisher.

**Supplementary Fig. 2 Sampling Strategy for Target Tissue Landmarks with Spatial Domain Associations (another example related to Fig. 3)**

This figure is similar to Fig. 3C-J. The difference is that the TTLs are simulated based on another spatial domain. A An example for cell type and TTL maps of a slice. B: Example of the cell type and the TTL maps in an FOV sampling sequence in color with unsampled part in grey. C: The Sampling frequency heatmap on a slice with w/ SOFisher within 1000 trails with randomly initial FOVs. D: The Sampling frequency heatmap on a slice with w/o SOFisher for performance comparison. E: The variation in the number of hits with the increase of steps for the two compared policies of w/ and w/o SOFisher. The p-values (denoted by P) between the hits of the two policies at steps 5, 10, 15, 20, 25, 30, 35, 40, 45, 50. F: The minimum number of FOVs for obtaining a target number (=10) of hits, and the number of the first hit in a sampling sequence for 1000 trails with randomly initial FOVs. The p-values of the two performance indices between w/ and w/o SOFisher are given. G: Comparison in terms of the number of hits between w/ and w/o SOFisher within 10, 20, 30, 40, 50 steps. H: In the table, the second and third lines show the average of the number of hits over 200 trails within each of 10, 20, 30, 40, 50 steps and over the total 1000 trails, and the last line show the improvements of w/ SOFisher over w/o SOFisher.

**Supplementary Fig. 3 The next sampling positions after executing an action (Related to Methods section)**

